# Rtt109 promotes nucleosome replacement ahead of the replicating fork

**DOI:** 10.1101/2022.02.09.479766

**Authors:** Felix Jonas, Gilad Yaakov, Naama Barkai

**Affiliations:** Department of Molecular Genetics, Weizmann Institute of Science, Israel

**Author notes:** Contributed equally (Felix Jonas, Gilad Yaakov).

## Abstract

DNA replication perturbs chromatin by triggering the eviction, replacement and incorporation of nucleosomes. How this dynamic is orchestrated in time and space is poorly understood. Here, we apply a recently established sensor for histone exchange to follow the time-resolved histone H3 exchange profile in budding yeast cells undergoing synchronous replication. We find that new histones are incorporated not only behind, but also ahead of the replication fork. We provide evidence that Rtt109, the S phase-specific acetyltransferase, stabilizes nucleosomes behind the fork, but promotes H3 replacement ahead of the fork. Unexpectedly, increased replacement ahead of the fork is independent of the primary Rtt109 acetylation target H3K56, and rather results from Rtt109 activity towards the H3 N-terminus. Our results suggest that selective incorporation of differentially modified H3s behind and ahead of the replication fork results in opposing effects on histone exchange, which may contribute to genome stability by overcoming replication-associated challenges.

## Introduction

Eukaryotic DNA is wrapped around histone octamers to form nucleosomes, the basic building blocks of chromatin (Luger et al. 1997). Nucleosomes limit access to the DNA, thus protecting the genome and providing a unified platform for regulating DNA-based processes (Kristjuhan and Svejstrup 2004; Schwabish and Struhl 2004; Bondarenko et al. 2006; Fleming et al. 2008; Lai and Pugh 2017). Regulatory mechanisms act on chromatin to change nucleosome positioning along the DNA, and to modify histone residues for controlling the recruitment of regulatory factors or histone-DNA affinity (Bannister and Kouzarides 2011; Owen-Hughes and Gkikopoulos 2012). Acetylation of histones on their N-terminus, for example, can reduce nucleosome stability and facilitate access to DNA (Hong et al. 1993; Lee et al. 1993; Bauer et al. 1994). Complementing these mechanisms that act on DNA-bound histones, is the process of nucleosome exchange where histone chaperones and remodelers evict or replace DNA-bound histones (Laskey et al. 1978; Dilworth and Dingwall 1988; Ito et al. 1996; Verreault et al. 1996; Andrews and Luger 2011; Hsieh et al. 2013; Mattiroli et al. 2015; Clapier et al. 2017; Hammond et al. 2017). Histone exchange reshapes the epigenetic landscape by diluting position-dependent marks while enriching for modifications present in the unbound histone pool.

Nucleosome dynamics are particularly prominent during DNA replication (Groth et al. 2007; Hamperl and Cimprich 2016). In addition to the increase of nucleosomes needed for wrapping newly synthesized DNA, progression of the replication fork requires nucleosomes to disassemble, at least transiently. Nucleosome disassembly is required at actively replicating locations, but nucleosome replacement could also extend to regions ahead of the fork, where accumulated helical tensions may evict nucleosomes (as shown during transcription (Corless and Gilbert 2016)) exposing DNA to potential damage. Finally, enzymes expressed in S phase modify histones prior to DNA incorporation, and these specifically modified histones become enriched on newly synthesized DNA, shaping an epigenetic landscape unique to replication with possible consequences on chromatin dynamics.

The S phase-specific acetyltransferase Rtt109 is a central regulator of the replication-specific epigenetic landscape in budding yeast. Rtt109 acetylates newly synthesized histones on their internal H3K56 residue, and on multiple N-terminal H3 residues (Han et al. 2007; Tsubota et al. 2007; Berndsen et al. 2008; Fillingham et al. 2008). The histone chaperone Asf1 is required for both H3K56 and H3 N-terminal acetylation by Rtt109, while another chaperone, Vps75, is required only for the latter (Berndsen et al. 2008; Fillingham et al. 2008). Of note, while both modifications are added prior to incorporation onto DNA, their genomic profile and temporal stability vary: H3K56 acetylation remains associated with replicated DNA until the end of S phase, when its specific deacetylases Hst3 and Hst4 are induced (Celic et al. ; Maas et al. 2006; Bar-Ziv et al. 2016; Voichek et al. 2016b; Voichek et al. 2018). By contrast, replication-dependent acetylation of H3K9, and likely other N-terminal residues, appears as a transient wave that closely associates, and in fact precedes, the replication fork (Bar-Ziv et al. 2016).

Functionally, cells are viable without Rtt109, but suffer from genomic instability (Driscoll et al. 2007; Li et al. 2008). The contribution of Rtt109 to additional cellular processes depends on its specific activity: H3K56 (but not H3K9) acetylation is required for expression homeostasis, namely the transcriptional buffering of the gene dosage increase in replicated regions (as compared to those not yet replicated) (Bar-Ziv et al. 2016; Voichek et al. 2016a; Voichek et al. 2016b; Voichek et al. 2018; Bar-Ziv et al. 2020). By contrast, Rtt109-dependent H3 N-terminal acetylation slows down the replication fork, a phenotype not seen in H3K56 mutants(Frenkel et al. 2021).

The fact that Rtt109 acts on unbound H3 suggests that it plays a role in H3 incorporation. Consistent with such a role, H3K56 acetylation increases the affinity of H3 towards CAF1, the histone chaperone incorporating histones at the replication fork (Li et al. 2008; Han et al. 2013). Genome-wide mapping of H3 exchange rates further revealed a tight correlation between histone exchange and H3K56 acetylation (Rufiange et al. 2007; Kaplan et al. 2008), and functional analysis of H3K56 mutants indicated a contribution of this modification to histone dissociation rates at particular locations (Ferrari and Strubin 2015) or *in-vitro* (Lee and Lee 2019). However, these studies correlating H3K56ac with histone exchange at the genomic level considered non-replicating cells, in which Rtt109 is repressed and H3K56ac levels are low (Rufiange et al. 2007; Kaplan et al. 2008). As for replication-associated H3K9ac, its localization ahead of the progressing fork (Bar-Ziv et al. 2016) may suggest a contribution to H3 incorporation there (Frenkel et al. 2021), but no functional data supporting this notion is available to date.

Clarifying the role of histone exchange in expression homeostasis and replication-fork slowdown requires measuring the nucleosome exchange pattern in replicating cells. Traditional techniques to assay exchange apply pulse-chase histone labeling followed by the collection of multiple time-resolved samples. These therefore involve temporal perturbations and are subject to significant time delays from histone labeling to exchange measurements, limiting their use during rapid dynamic processes such as DNA replication (Ahmad and Henikoff 2002; Keppler et al. 2004; Schermer et al. 2005; Dion et al. 2007; Rufiange et al. 2007; Kaplan et al. 2008; Verzijlbergen et al. 2010; Radman-Livaja et al. 2011; Verzijlbergen et al. 2011; Smolle et al. 2012; Venkatesh et al. 2012; Das and Tyler 2013; Huang et al. 2013; Kraushaar et al. 2013; Ha et al. 2014; Yildirim et al. 2014; Sadeghi et al. 2015; Siwek et al. 2018). Recently, we developed a new histone exchange sensor that alleviates the need for pulse-chase by using genetically encoded exchange sensors, and allows for exchange measurements during a dynamic process using a single sample (Yaakov et al. 2021). Here, we use wild type and mutant strains carrying an H3 exchange sensor to define the time-resolved, genome-wide H3 exchange pattern during slowed replication. We further test the roles of Rtt109 and its associated H3 acetylations in regulating this exchange during replication.

## Results

### The H3 exchange profile during replication

We engineered the H3 exchange reporter strain as described and validated in (Yaakov et al. 2021) by fusing both H3 alleles to a TEV-cleavable tag and H2B to the TEV protease (Methods). The H3 tag is composed of two epitopes (HA and myc) connected via a linker with a TEV cleavage site. Since H3 and H2B bind exclusively in the context of the histone octamer, myc cleavage only occurs after nucleosome formation on DNA. Accordingly, at any given genomic locus, HA levels report on nucleosome (H3) occupancy, myc levels report on the associated H3 incorporation rate, and their ratio on the exchange rate(Yaakov et al. 2021).

To enable sufficient temporal resolution, we slowed replication by subjecting cells to the deoxynucleotide-depleting drug hydroxyurea (HU). Cells carrying the H3 exchange reporter were G1-arrested, released into HU-containing medium, and sampled every 10-20 minutes up to 3 hours (Figure 1A). ChIP-seq was used to profile the H3 exchange tags, HA and myc, genome-wide. Using the same samples, we also mapped two Rtt109-dependent modifications, H3K56 and H3K9 acetylation.

**Figure 1.**
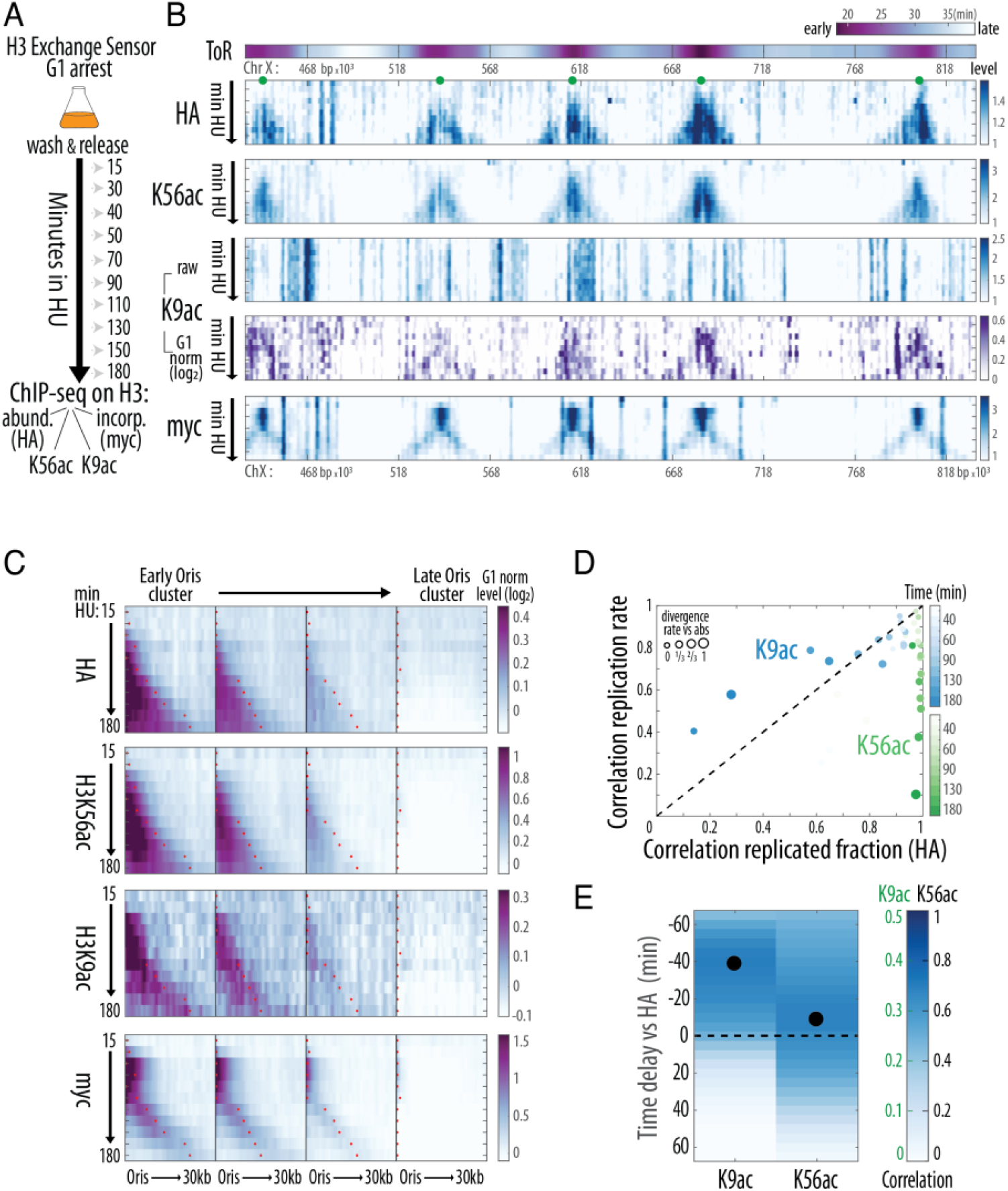
Chromatin replication dynamics during slowed S phase. A, B) *Profiling histone H3 modifications and exchange dynamics during slowed replication:* Time-resolved Chromatin ImmunoPrecipitation (ChIP-seq) experiments were performed to follow replication in cells synchronously released into hydroxyurea (HU) to slow replication. H3 abundance (HA), H3 incorporation (myc, see (Yaakov *et al*., 2021)) and H3K56 & H3K9 acetylations were measured (A). Replication dynamics are captured by the change in read coverage, shown here on a segment of Chromosome X (B). The annotated Time of Replication (ToR) is included (Yabuki et al., 2002). Annotated origins of replication initiation (ORIs, (Siow et al., 2011)) with ToR<21 minutes are show as green dots. H3K9ac coverage is additionally shown after normalizing to non-replicating, G1-arrested cells (log_2_). C, D) *H3K56ac correlates with DNA abundance, while H3K9ac accumulates ahead of the replication fork:* ORIs were clustered by their ToR, and profiles around each ORI were aligned and averaged. Shown are G1-normalized profiles within each of the four ORI clusters (C, note the scales for the various antibodies) and the fork position was calculated from the HA profile (red dots). Data for H3K9ac or H3K56ac (n=2 time courses) are summarized (D) by comparing the profiles of these modifications with the HA profile (replicated fraction) or its change along the genome (replication rate, see Methods). Color intensity indicates time, and the dot size indicates the divergence (1-correlation) between the replicated fraction and rate. E) *H3K9ac accumulates ahead of the replication fork:* Cross-correlation between the temporal changes of the indicated acetylation with HA is shown as a function of the time-delay, black dots indicate the delay with the highest correlation (see Figure S1 for repeat). H3K56ac coincides with HA (∼DNA content), while H3K9ac precedes it. Note the different color scales for the epitopes.

Histone exchange is composed of replication-dependent and independent components. The former localizes to regions surrounding the fork, while the latter also occurs in regions not actively replicating (associated e.g. with transcription). To distinguish these contributions, we defined the temporal progression of DNA replication, using nucleosome abundance (HA) as an internal indicator of DNA content. As expected, nucleosome abundance increased gradually, with different genomic regions replicating at different times, in tight correlation with their annotated Time of Replication (ToR) (Figure 1B). Using this analysis, we defined for each time point the fraction of cells in which a given region has already been replicated (replicated fraction), and the fraction of cells in which replication is active within the current timeframe (replication rate, Methods).

Examining the Rtt109-dependent marks verified the expected localization of H3K56ac to replicated regions (Figure 1B,C). The H3K9ac profile, on the other hand, is composed of transcription and replication-dependent components, with only the latter being Rtt109-dependent (Bar-Ziv et al. 2016), and therefore requires normalization by the H3K9ac profile of non-replicating (G1-arrested) cells (Figure 1B). Normalized H3K9ac localized to regions that were actively replicating, but contrasting H3K56ac, did not accumulate in replicated regions (Figure 1C,D). Examining the temporal relation of the H3K9ac profile to the replication fork verified that, similar to that of rapidly replicating cells (Bar-Ziv et al. 2016), H3K9ac precedes the progressing fork in our slowed replication time course (Figures 1E & S1A).

To quantify H3 exchange, we considered the myc profile reporting on the H3 incorporation rate. As expected, once replication commenced, myc began to increase around early-replicating origins and then expanded and spread to later replicating regions (Figure 1B,C). Subsequently, the myc signal decreased in replicated regions, while still localizing to actively replicating ones (Figure 1B,C). Reduction of myc in replicated regions was partial, as it remained higher than at loci not yet replicated (Figure S1B). Finally, high myc signal was also found ahead of the replication fork, similar to H3K9ac (Figures 2A & S1A). We conclude that the myc sensor successfully captures histone incorporation during the dynamic process of replication.

**Figure 2.**
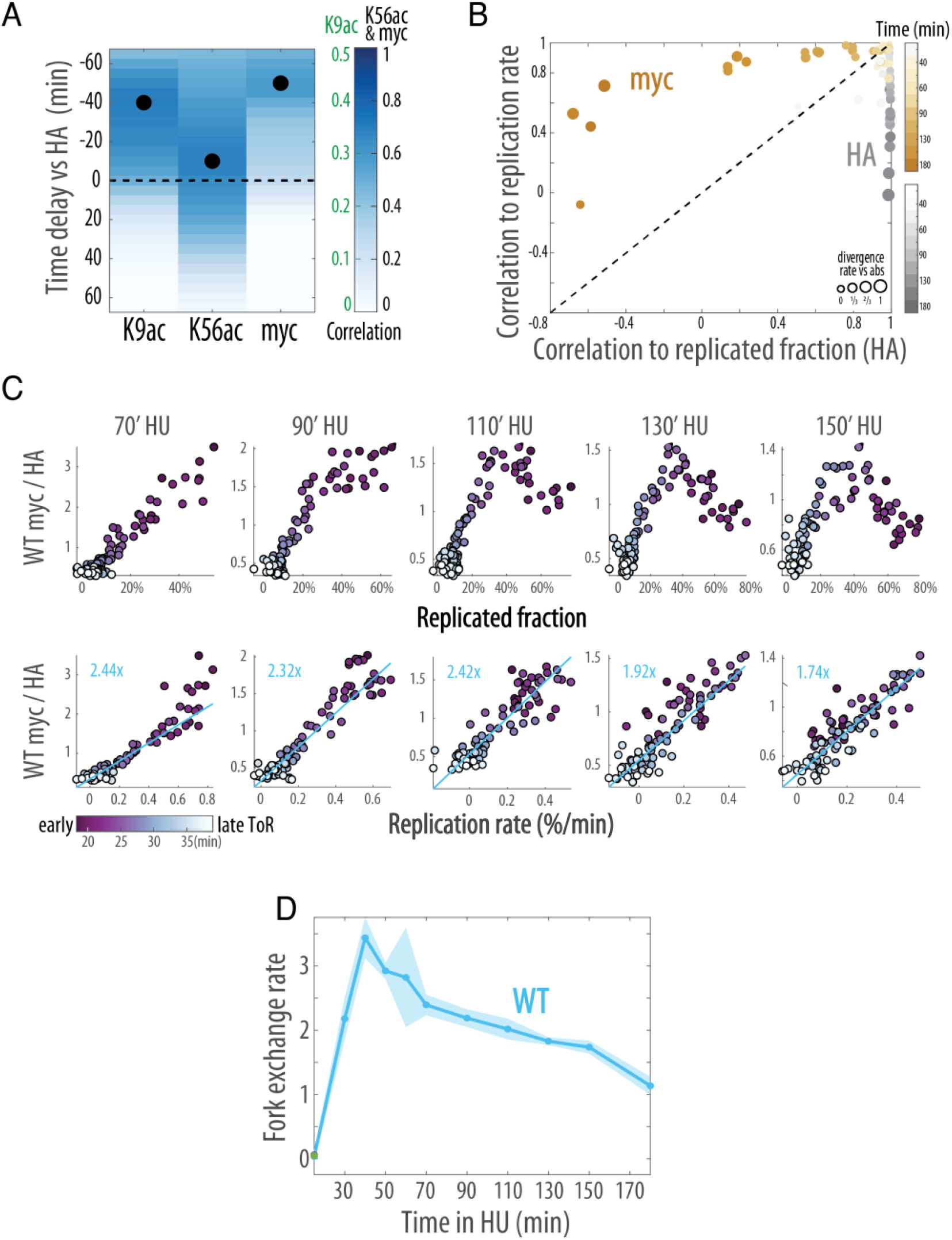
H3 exchange follows replication fork progression. A, B) *Histone incorporation precedes the replication fork and correlates with the replication rate*. (A) As figure 1E, adding the myc epitope of the exchange sensor marking new histones. Note the different color scales for the epitopes. See Figure S1 for additional repeats (n= 4 time courses). (B) myc correlation to replication fraction and rate, analyzed as Figure 1D. C, D) *Scaling of histone exchange with replicated fraction and replication rate*. The genome was clustered based on the annotated ToR, and the indicated profile was averaged within each of the 96 clusters (median). (C) shows the average H3 exchange (myc/HA) as a function of the HA-derived replicated fraction (top) or replication rate (bottom). Clusters are colored by their respective ToR. Time points 70-150 of the time course are shown (see Figure S2 for all time points). The scaling of exchange with the replication rate is summarized in (D), displaying the slopes of the linear fits in (C) (blue lines, Figure S2). Shading indicates standard error (n=4).

### Replication rates dominate the replication-dependent H3 exchange

Different processes contribute to replication-dependent histone exchange. These include the perturbation of nucleosomes ahead of the fork and the incorporation of new histones needed for wrapping the newly synthesized DNA. In addition, nucleosome dynamics may change due to the unique epigenetic landscape generated through replication, e.g. enrichment of H3K56ac. As replication is stochastic, these three processes overlap spatially and temporally, even in our synchronized cultures. In particular, regions of increased gene dosage (replicated in some cells) are still being actively replicated in other cells.

To distinguish the contribution of these individual processes to the myc profile, we compared H3 exchange (myc/HA) at each genomic region to both DNA content (the replicated fraction) and to the rate at which this DNA content increases (replication rate) (Figures 2B,C & S2A,B). Throughout our time course, H3 exchange remained tightly correlated with the replication rate, but not with the DNA content (Figures 2B,C & S2C). Quantitatively, the scaling of H3 exchange at the fork with the replication rate stayed relatively constant throughout the time course (Figure 2D). We conclude that in wild type cells, nucleosome incorporation in proximity to the progressing replication fork dominates the nucleosome exchange pattern.

In line with previous studies (Rufiange et al. 2007; Kaplan et al. 2008), we detected a clear correlation between H3K56ac and H3 exchange in non-replicated cells (Figure S2C), and accordingly expected to observe higher exchange in replicated regions where K56ac accumulates. We noted, however, that the correlation was lost during replication. In particular, H3K56ac was stably maintained after replication concluded, while myc was mostly lost (Figures 1B,C & S1B).

### Rtt109 deletion reduces nucleosome incorporation at the fork

Our results suggest that H3K56ac does not promote nucleosome exchange, at least during S phase. Since these results are at odds with previous suggestions, we wished to examine the role of H3K56ac in nucleosome exchange more directly. For this, we repeated the experiment in cells deleted of Rtt109 (Figure S3A,B), the only enzyme that deposits this modification. Rtt109 deletion was of little consequence on nucleosome exchange in G1-arrested cells and at early time points after release (Figures 3A and S3C). The exchange pattern did change, however, once replication progressed: it lost its tight correlation with replication rate, and instead became correlated with the replicated fraction (Figure 3A,B).

**Figure 3.**
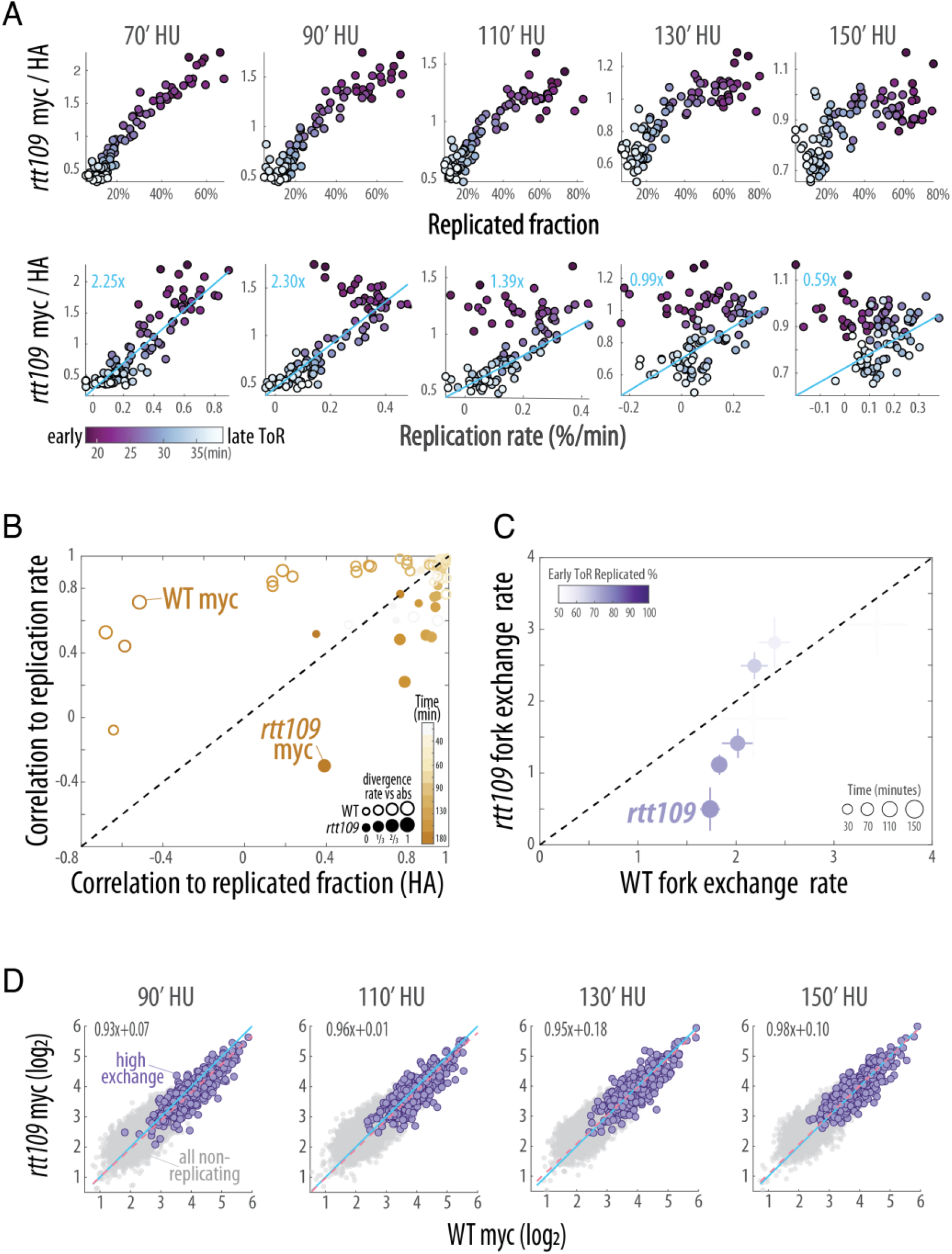
Scaling of H3 exchange with the replication rate depends on Rtt109. A-B) *Exchange in RTT109-deleted cells is dominated by the replicated fraction, rather than the replication rate:* (A) and (B) as Figure 2C and 2B, respectively. See Figure S3 for all time points. C) *Exchange at the replication fork is decreased in RTT109-deleted cells*. The average fork exchange rate in RTT109-deleted cells is plotted against the corresponding rate in wild type cells. Error bars indicate the standard error (n=3 for *rtt109*, n=4 for WT). Color intensity indicates the replicated fraction at loci with the earliest ToR, and dot size indicates time. D) *Replication-independent histone dynamics are invariant to RTT109 deletion*. Mean myc level of nucleosomes that have not yet been replicated at the indicated time points in wild type versus RTT109-deleted cells (n=4 and 3 respectively). Highlighted are the rapidly exchanging nucleosomes (as determined in G1-arrest) used for the linear fit (pink dotted line and equation). The 1-to-1 line is shown in blue.

This became evident at the later time points, when the replication of early replicating regions approached completion. Of note, the replication pattern, as captured by nucleosome occupancy (HA), remained largely invariant to Rtt109 deletion (Figure S3A). Therefore, in Rtt109-deleted cells, H3 exchange signal is reduced at the replication fork, while increasing in replicated regions (Figure 3A,C & S3E).

Since our data is relative, reduced myc levels at the fork could either indicate reduced incorporation at these regions, or increased incorporation elsewhere. Refuting the latter possibility, myc levels at rapidly exchanging nucleosomes that have not yet been replicated remained invariant to Rtt109 deletion (Figure 3D). Wild type cells therefore show more exchange per replicated nucleosome as compared to Rtt109-deleted cells. We conclude that for a given replicated nucleosome, wild type cells replace multiple nucleosomes at the fork or in its vicinity, and that these multiple replacements depend on Rtt109.

### Rtt109-dependent H3 N-terminal acetylation is required for fork-associated H3 replacement

Rtt109 thus contributes to replication-associated nucleosome exchange in two ways. First, by stabilizing nucleosomes in replicated regions, and second by increasing nucleosome replacement associated with the progressing fork. As shown above, the two types of Rtt109 acetylation differentially localize to either replicated regions (H3K56ac) or ahead of the progressing fork (H3K9ac). Considering this pattern, we hypothesized that Rtt109 promotes nucleosome replacement ahead of the fork by acetylating the H3 N-terminus, while contributing to nucleosome stability in replicated regions through H3K56 acetylation.

To examine this hypothesis, we repeated the experiment in cells deleted of Vps75 (Figures 4A & S4A), a chaperone that cooperates with Rtt109 in acetylating the H3 N-terminus, but is dispensable for H3K56 acetylation. Consistent with this role of Vps75, its deletion largely abrogated the replication-associated H3K9ac localization ahead of the fork (Figures 4B & S4A-C), while having no effect on H3K56ac (Fig S4A,B, 4B). Examining the exchange pattern confirmed that Vps75 deletion specifically reduced fork-associated exchange, resembling the loss of correlation observed in Rtt109-deleted cells (Figures 4C & S4D-F). As further predicted, Vps75 deletion had little effect on the exchange at replicated and non-replicated regions (Figure 4A, S4F). Together, we conclude that by acetylating the H3 N-terminus, Rtt109 promotes nucleosome replacement ahead of the progressing fork, but H3K9ac does not affect nucleosome exchange in replicated regions.

**Figure 4.**
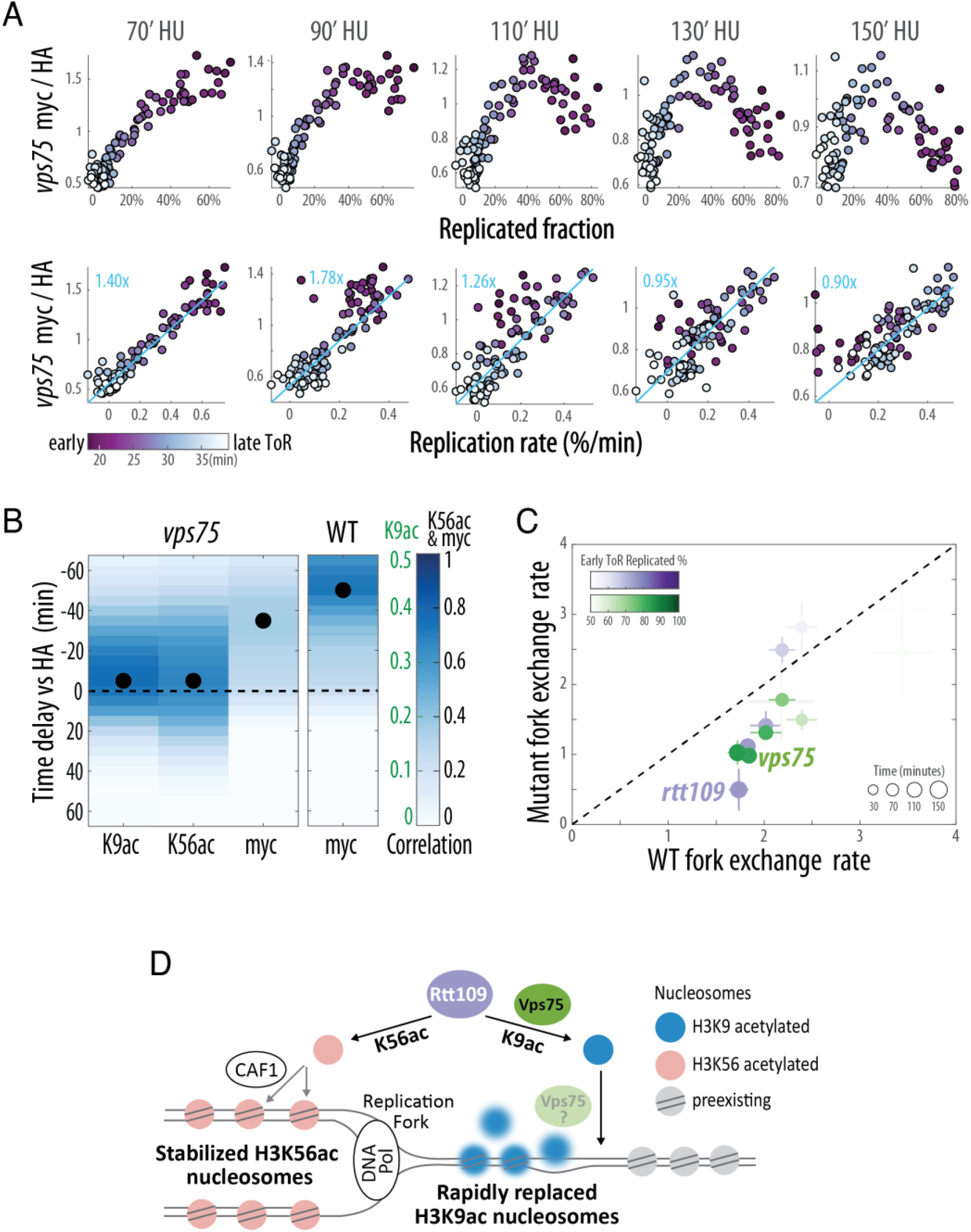
N-terminal H3 acetylation mediates histone exchange ahead of the fork. A-C) *VPS75-deleted cells lose replication-associated K9ac and display reduced H3 exchange ahead the fork, but stable nucleosomes behind it*. (A-B) as Figure 2C and 2A respectively for VPS75-deleted cells (n=2, see also Figure S4). Of note is the delayed H3K9 acetylation (B) (compare to wild type cells Figures 2A & S1A). Note the different color scales for the epitopes, WT myc correlation is shown for comparison. (C) as Figure 3C with the addition of the *vps75* mutant. D) *Rtt109’s dual role in regulating histone exchange during DNA replication*. A schematic model describing the main findings: Rtt109-dependent acetylation of K56 stabilizes nucleosomes behind the replication fork, while acetylation of K9 promotes nucleosome replacement ahead of the fork. See Discussion.

## Discussion

The packaging of eukaryotic genomes within chromatin poses a major obstacle for DNA replication, necessitating the rearrangement of chromatin. In this study, we describe a dual role of Rtt109 in orchestrating nucleosome dynamics during DNA replication, implemented through its distinct activities towards H3K56 and the H3 N-terminus. First, by acetylating H3K56, Rtt109 stabilizes nucleosomes at regions already replicated. Second, by acetylating the H3 N-terminus, Rtt109 promotes H3 replacement ahead of the replication fork (Figure 4D).

The acetylation of H3K56 increases its affinity to CAF1, the histone chaperone that incorporates nucleosomes in the wake of the fork (Li et al. 2008; Han et al. 2013). H3K56ac then accumulates in replicated regions where it remains throughout S phase. As studies had implicated the H3K56ac modification in promoting the rate of nucleosome exchange (Rufiange et al. 2007; Kaplan et al. 2008; Ferrari and Strubin 2015), we previously suggested that this continuous presence of H3K56ac results from rapid H3 exchange in these regions, thus replenishing H3K56ac from the enriched unbound pool(Voichek et al. 2018). This was supported by a reduced accumulation of position-specific marks through e.g. gene expression, which we previously reported and suggested to contribute to H3K56ac-dependend expression homeostasis (Voichek et al. 2018). Contradicting this hypothesis, our exchange reporter (myc) did disappear from replicated regions that still displayed high H3K56ac. Moreover, deletion of Rtt109 led to a higher, rather than lower exchange rate in replicated regions. This increased exchange in replicated regions was absent in *vps75* mutant cells, confirming that H3K56ac was sufficient for stabilizing replicated nucleosomes. Therefore, during S phase, H3K56ac acts to stabilize, rather than destabilize replicated nucleosomes. Whether and how this increased stability contributes to expression homeostasis remains to be studied.

Contrasting H3K56ac, which contributes to expression homeostasis but has no effect on fork velocity, we recently showed that Rtt109-dependent acetylation of the H3 N-terminus, which is dispensable for expression homeostasis, slows down replication (Frenkel et al. 2021). Since Rtt109-dependent H3K9ac localizes ahead of the fork, we suggested that this slowdown results from increased nucleosome replacement ahead of the fork (Frenkel et al. 2021). We now provide direct evidence supporting this model: the rapid nucleosome replacement observed in front of the fork is suppressed by deletion of Rtt109 and, more specifically by deletion of Vps75, linking it directly to N-terminal H3 acetylation.

We initially hypothesized that nucleosome replacement ahead of the fork is triggered by the helical stresses that accumulate ahead of the fork, and that acetylated H3 is preferentially incorporated in these nucleosome-lacking regions to protect the exposed DNA (Frenkel et al. 2021). However, we could not detect a decrease in nucleosome abundance ahead of the fork in neither *rtt109* nor *vps75* mutants. Neither could we detect expansion of H3K56ac pattern upon Vps75 deletion, expected in such a scenario. We therefore find it more consistent that Vps75 directly replaces nucleosomes ahead of the fork, thereby preparing the chromatin for the approaching fork (Figure 4D).

Rtt109 is not essential, but is required for genome stability (Driscoll et al. 2007; Li et al. 2008). Controlled histone exchange may be the molecular mediator of this function. In replicated regions, increased nucleosome stability is evoked by H3K56 acetylation. At the same time, increased exchange ahead of the fork functions to maintain fork velocity, which could also be important for genomic stability. Indeed, perturbations that either reduced this velocity, like nucleotide depletion (Bester et al. 2011), or increased it, for example through PARP inhibition (Maya-Mendoza et al. 2018), were found to reduce genome stability. Therefore, Rtt109, through its two distinct functions, regulates nucleosome exchange in apparently opposing ways, compatible with the respective stresses challenging genome stability.

---

**Figure S1.**
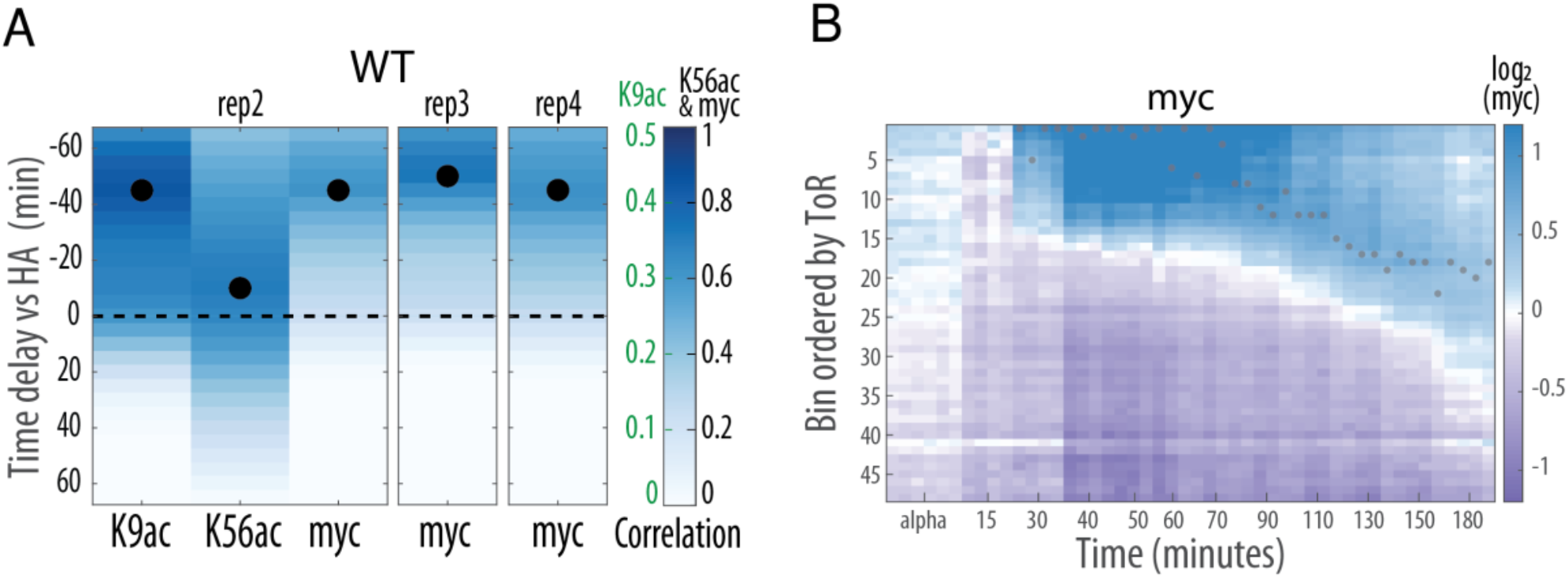
Myc and acetylation dynamics in wild type cells. A) *Temporal sequence of Rtt109-dependent histone marks and H3 integration during DNA replication*. Cross-correlation analysis as in Figure 1E. Additional repeats of the time course experiment consistently measure time delays between the various epitopes, note the different color scales for the epitopes. B) *Myc dynamics in replicated versus non-replicated regions*. Median myc coverage in ToR clusters (Y-axis) as a function of experiment time point in all time courses (X-axis, each column is a time course centered around the time mark, n=4). Grey dots indicate HA-derived position of the replication fork.

**Figure S2.**
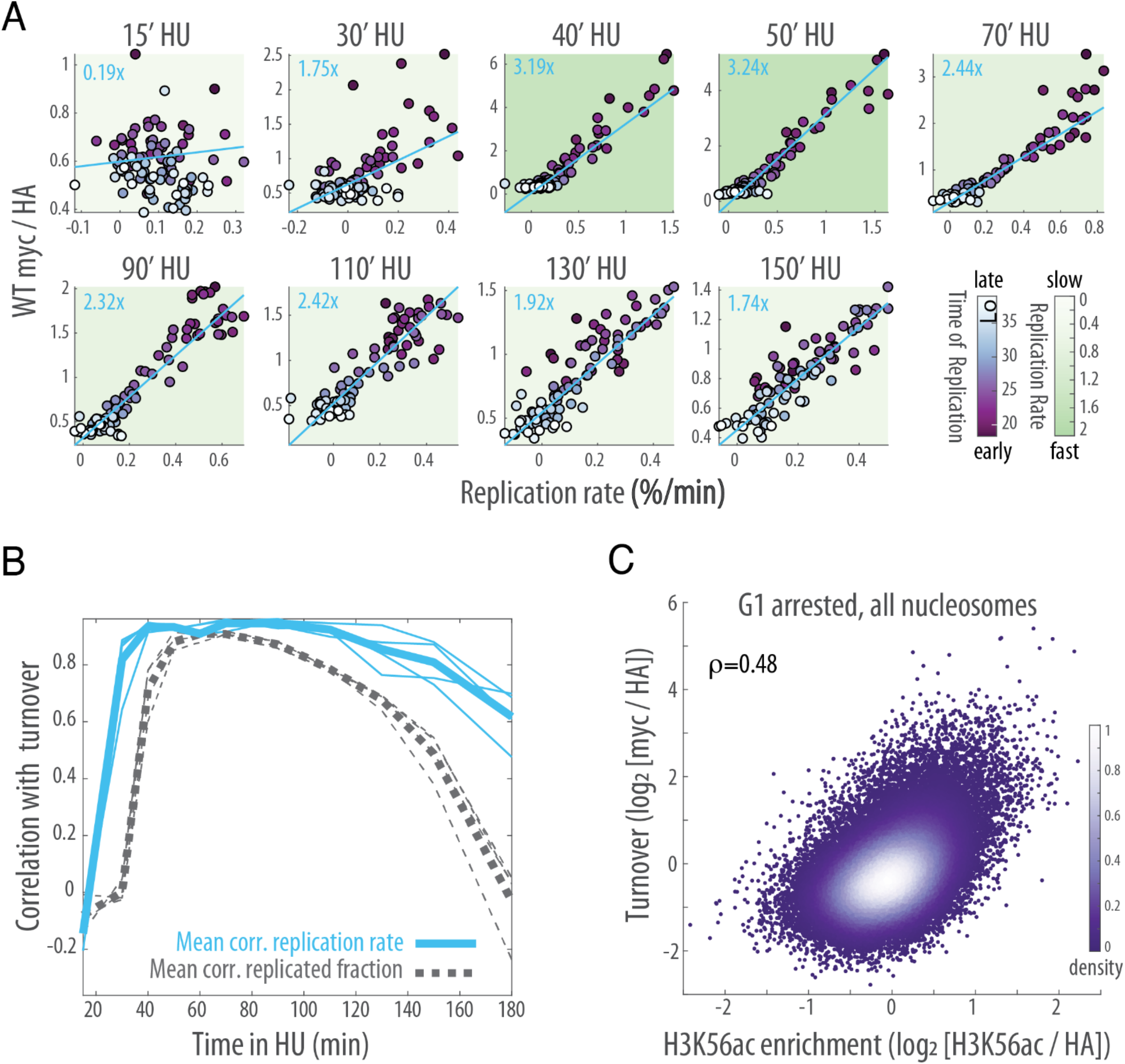
H3 exchange during DNA replication. A-B) *H3 exchange is strongly correlated with replication rate throughout the time course*. (A) as Figure 2A for all time points, with the background color indicating the maximal replication rate at a given timepoint. The linear fit and its slope measuring fork exchange is shown in blue. (B) Correlation between H3 turnover (log_2_(myc/HA)) and replication rate or replicated fraction across all clusters. The thick line corresponds to the mean of 4 experiments indicated in thin lines. Note the drop in correlation at the later times. C) *Turnover correlates with H3K56ac in G1-arrested cells*. Histone turnover (log_2_ myc/HA) and acetylation enrichment (log_2_(K56ac /HA)) of wild type cells in non-replicating G1-arrested cells.

**Figure S3.**
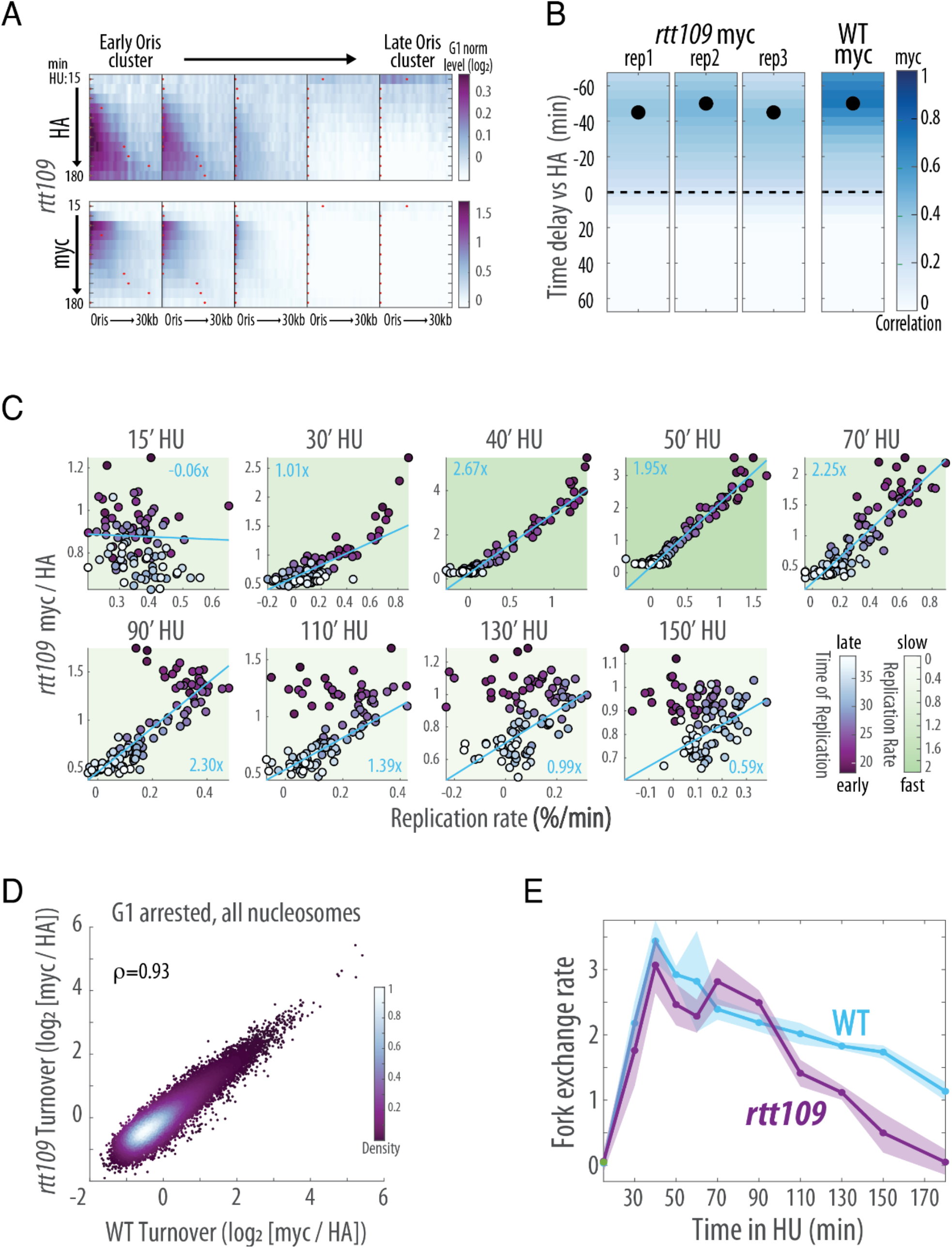
Replication-associated histone dynamics in RTT109-deleted cells. A) *H3 sensor levels around ORIs of RTT109-deleted cells*. As Figure 1C for RTT109-deleted cells with same scale intensities. B) *Temporal relation between myc and fork progression in RTT109-deleted cells*. As Figure S1A for three repeats of RTT109-deleted cells. WT myc correlation is shown comparison. C) *Loss of correlation between H3 exchange and replication rate at mostly replicated regions after RTT109 deletion*. As Figure S2A for RTT109-deleted cells. Note the divergence of early replicating clusters from the line in later time points when they are mostly (>70%) replicated. D) *Replication-independent histone dynamics are invariant to RTT109 deletion*. Mean turnover (log_2_ myc/HA) of *rtt109* versus WT cells showing all nucleosomes (n=5 for each strain). E) *Reduced fork-associated histone exchange in RTT109-deleted cells*. As Figure 2D for the indicated strains (n=4 for WT, n=3 for *rtt109*). Shading is standard error.

**Figure S4.**
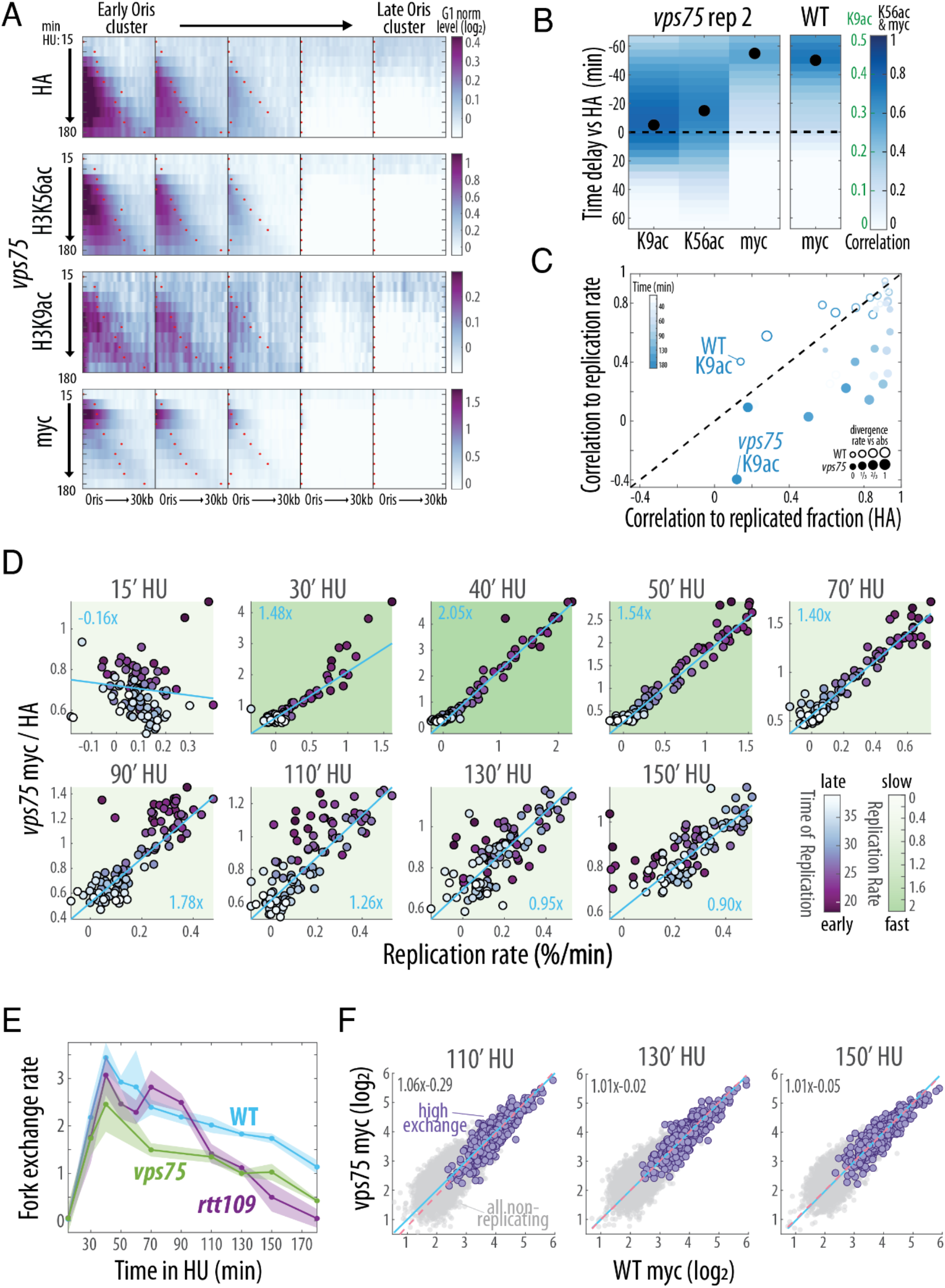
Replication-associated histone dynamics in VPS75-deleted cells. A-C) *VPS75 deletion removes fork associated K9 acetylation*. (A-C) as Figures 1C, 1E and 1D respectively for the indicated epitopes in the *vps75* mutant. (B) is a repeat for Figure 4B, alongside myc WT correlation for comparison. (C) Both *vps75* repeats are included. D-E) *Reduced fork exchange rate in VPS75-deleted cells*. (D) as Figure S2A for VPS75-deleted cells. (E) as Figure S3E with the addition of the *vps75* strain (n=2). Shaded line is standard error. F) *Replication-independent histone dynamics remain unaffected by Vps75 deletion*. As Figure 3D for VPS75-deleted cells (mean of n=2).

**Figure S5.**
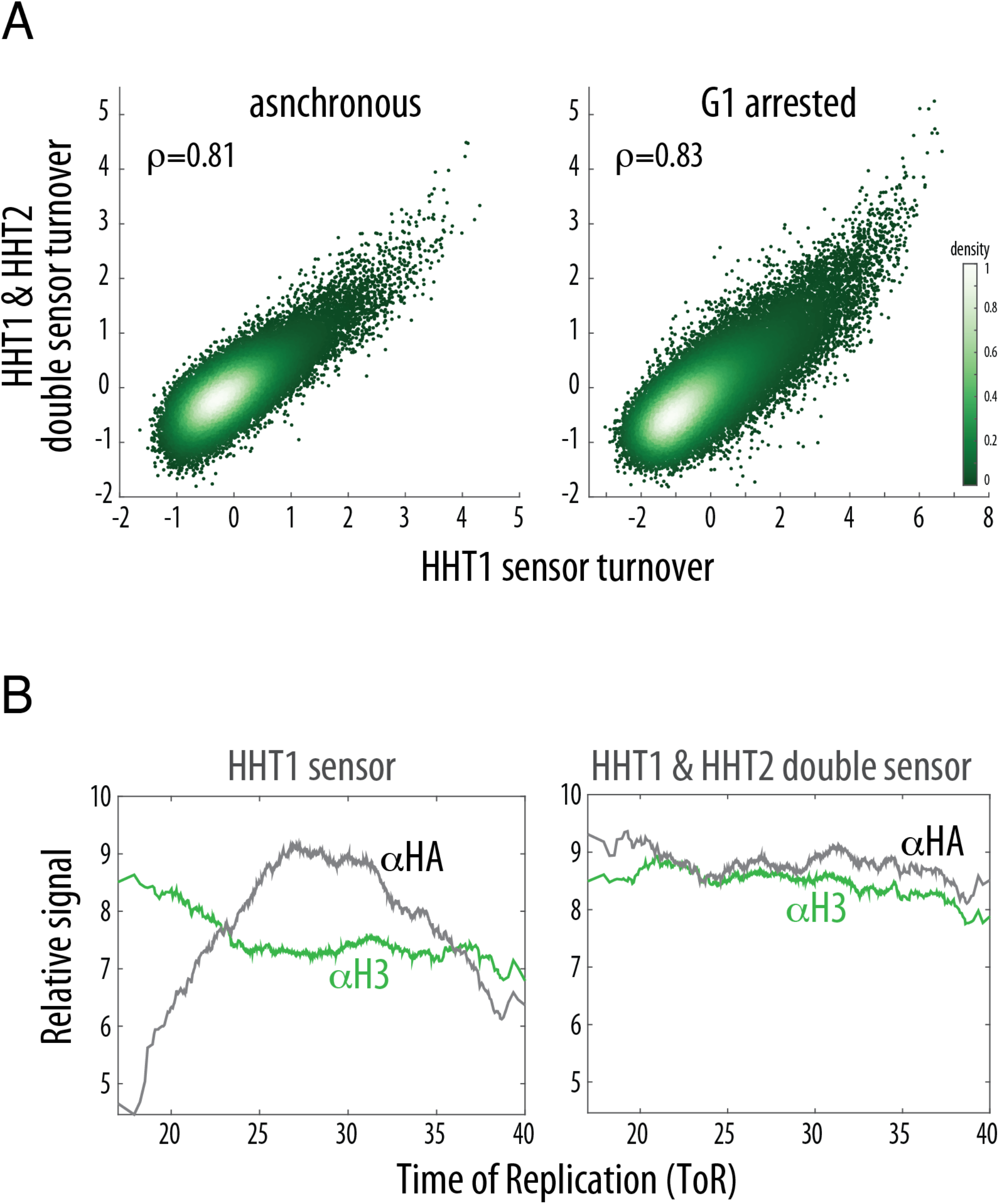
An H3 sensor to measure nucleosome exchange during DNA replication. A-C) *Strains with both H3 alleles tagged with the exchange sensor accurately measure replication-independent histone turnover, and ameliorate replication-associated allele imbalances*. (A) Turnover (log_2_(myc/HA)) of all nucleosomes in asynchronously growing (left) or non-replicating G1-arrested (right) cells with the single-tagged (HHT1) versus double-tagged (HHT1 and HHT2) H3 sensors. (B) Mean nucleosome occupancy of total histone H3 (using an anti-H3 antibody) versus sensor-tagged histone H3 (anti-HA antibody) across all ToRs. Note the replication-associated imbalance of the HHT1-tagged single-allele sensor that is abolished in the double-tagged sensor.

## Methods

### Yeast strains

All strains were generated from YGY663 (Yaakov et al. 2021) using standard yeast transformation (Gietz and Schiestl 2007). To prevent competition between tagged and non-tagged histone variants, we fused the HA-TEVsite-MYC sensor to Hht2 in YGY663 (in which only Hht1 is tagged), resulting in the H3 double-tagged replicate strains YGY672 and YGY673. We then confirmed almost identical turnover profiles and removal of competition in growing and G1-arrested yeast cells (see Figures S5A-B). Rtt109 (YGY674 & YGY675) or Vps75 (YGY721 & YGY722) were deleted using natNT2 or hphNT1 selection markers respectively (Janke et al. 2004) (see Table below).

Histone fusions and BAR1 deletion were introduced using CRISPR/Cas9 as in (Yaakov et al. 2021). Sequences including 200 bp up and downstream of the modified histone fusions are provided below. Silent mutations that introduce a mismatch/es to rescue from the guide RNA target sequence are indicated in blue. TEV carries an S219P mutation to inhibit (Yi et al. 2013). 2x(GGSG) linkers were added between the C-terminus of each modified histone subunit and its fusion. Note that the 5’ and 3’ UTRs regions in these modified loci remain untouched, such that the fusion is under the control of the native promoter and terminator. All other histone genes remain wild type.

### HTB2-TEV

ataaggttttggatcagtaaccgttatttgagcataacacaggtttttaaatatattattatatatcatggtatatgtgtaaaatttttttgctgactggttttgtttatttatttagctttttaaaaattttactttcttcttgttaattttttctgattgctctatactcaaaccaacaacaacttactctacaactaatgtcctctgccgccgaaaagaaaccagcttccaaagctccagctgaaaagaagccagctgccaagaaaacatcaacctccgtcgatggtaagaagagatctaaggttagaaaggagacctattcctcttatatttacaaagttttgaagcaaactcacccagacactggtatttcccagaagtctatgtctattttgaactctttcgttaacgatatctttgaaagaattgctactgaagcttctaaattggccgcttataacaagaaatccactatttctgctagagaaatccaaacagccgttagattgatcttacctggtgaattggctaaacatgccgtctccgaaggtactagggctgttaccaaatactcctcatctactcaagccggaggtagtggagggggatcaggaaaaagcttgtttaaggggccgcgtgattacaacccgatatcgagcaccatttgtcatttgacgaatgaatctgatgggcacacaacatcgttgtatggtattggatttggtcccttcatcattacaaacaagcacttgtttagaagaaataatggaacactgttggtccaatcactacatggtgtattcaaggtcaagaacaccacgactttgcaacaacacctcattgatgggagggacatgataattattcgcatgcctaaggatttcccaccatttcctcaaaagctgaaatttagagagccacaaagggaagagcgcatatgtcttgtgacaaccaacttccaaactaagagcatgtctagcatggtgtcagacactagttgcacattcccttcatctgatggcatattctggaagcattggattcaaaccaaggatgggcagtgtggcagtccattagtatcaactagagatgggttcattgttggtatacactcagcatcgaatttcaccaacacaaacaattatttcacaagcgtgccgaaaaacttcatggaattgttgacaaatcaggaggcgcagcagtgggttagtggttggcgattaaatgctgactcagtattgtgggggggccataaagttttcatgccgaaacctgaagagccttttcagccagttaaggaagcgactcaactcatgaattaagtcactcactaggtattgtgatttagtcatgttttctttttattagtggcattttttatatattgtaacattagggttctgttacttgtccagatatgattggaagggtctagtctatcagcctccgaagggagttgtataaatgtatatatataactttataaacaaaaaaaattgggaaagatattttgcaaacagtt

### HHT1-HA-TEVsite-MYC

tatctctccttgacttttagcgtggaagataacgaaatgcccgggcgatttttctttttggtaccctccacggctccttgttgaaatacatatataaaagactgtgtattcttcgggatacatctctttcctcaaccttttatattctttctttctagttaataagaaaaacatctaacataaatatataaacgcaaacaatggccagaacaaagcaaacagcaagaaagtccactggtggtaaggccccaagaaagcaattagctt ctaaggctgccagaaaatccgccccatctaccggtggtgttaagaagcctcacagatataagccaggtactgttgctttgagagaaatcagaagattccaaaaatctactgaactgttgatcagaaagttgcctttccaaagattggtcagagaaatcgctcaagatttcaagaccgacttgagatttcaatcttctgccatcggtgccttgcaagaatctgtcgaagcctacttagtctctttatttgaagataccaacttggctgccattcacgccaagcgtgtcactatccaaaagaaggatatcaaattggccagaagactaagaggtgaaagatcaggaggtagtggcgggggaagtggatatccttatgatgtgcctgactacgctgggtatccttacgacgttccggattacgccggctcctacccgtatgacgtcccggactatgctactgaaaatctttatttccagtcaggtacgcgtaggtgggccagtggtgaacaaaaactgatatcagaggaagacctaaacggggaacaaaagcttataagcgaggaagatttaaatggagagcagaagcttatcagcgaagaagaccttaatggaagctcccgtggggagcaaaagctgatttctgaagaggatttgaatggagagcaaaaactgatttccgaagaggacctgaatggggagcaaaaactaatatcagaggaggatctgaacgggtcctcagaacagaagttgataagtgaggaagatttaaacggtgaacaaaaattgattagtgaggaggacttgtagtttgttgattgtcatcagttttagtaaaaaacgaacaaaaacacaataaaatataaatcaatatatttaggtttactgggttctttaacagttgtataatagttattttttattacaaaaatataggttttaataaaaaaaaatagggttctatttgttttacatttattgatttgtttttcctggcgataccctcgaaa

### HHT2-HA-TEVsite-MYC

acggctatggctcggtgtcaaaacatagtttgcgtgataacagcgtgttgtgctctctcgcgttgcttcttgtgaccgcagttgtatataaataatctttttcttgttcttttatataggaccactgttttgtgacttccactttggcccttccaactgttcttccccttttactaaaggatccaagcaaacactccacaatggccagaactaaacaaacagctagaaaatccactggtggtaaagccccaagaaaacaattagcctccaaggctgccagaaaatccgccccatctaccggtggtgttaagaagcctcacagatataagccaggtactgttgccttgagagaaattagaagattccaaaaatctactgaactgttgatcagaaagttacctttccaaagattggtcagagaaatcgctcaagatttcaagaccgacttgagatttcaatcttctgctatcggtgctttgcaagaatccgtcgaagcatacttagtctctttgtttgaagacactaatctggctgctattcacgctaagcgtgttactatccaaaagaaggatatcaaattggccagaagactaagaggtgaaagatcaggaggtagtggcgggggaagtggatatccttatgatgtgcctgactacgctgggtatccttacgacgttccggattacgccggctcctacccgtatgacgtcccggactatgctactgaaaatctttatttccagtcaggtacgcgtaggtgggccagtggtgaacaaaaactgatatcagaggaagacctaaacggggaacaaaagcttataagcgaggaagatttaaatggagagcagaagcttatcagcgaagaagaccttaatggaagctcccgtggggagcaaaagctgatttctgaagaggatttgaatggagagcaaaaactgatttccgaagaggacctgaatggggagcaaaaactaatatcagaggaggatctgaacgggtcctcagaacagaagttgataagtgaggaagatttaaacggtgaacaaaaattgattagtgaggaggacttgtgaatataaagcgggaatttttttttttctatgcatttagactggggggacatcatacaaacatctccttttttaatacattattcatctatctattcgcaatgcttttaacgacatgaggagggtatatatgctgtaatttatcgtttacagacataattgcggggctaatttttagatgcttagatagtatacgtaatttt

|  |  |
| --- | --- |
| YGY663 | MATa his3Δ1 leu2Δ0 LYS2 met15Δ0 ura3Δ0 HTB2-TEV, HHT1-HA-TEVsite-MYC, *bar1* |
| YGY672 | MATa his3Δ1 leu2Δ0 LYS2 met15Δ0 ura3Δ0 HTB2-TEV, HHT1-HA-TEVsite-MYC, HHT2-HA-TEVsite-MYC, *bar1* |
| YGY673 | MATa his3Δ1 leu2Δ0 LYS2 met15Δ0 ura3Δ0 HTB2-TEV, HHT1-HA-TEVsite-MYC, HHT2-HA-TEVsite-MYC, *bar1* |
| YGY674 | MATa his3Δ1 leu2Δ0 LYS2 met15Δ0 ura3Δ0 HTB2-TEV, HHT1-HA-TEVsite-MYC, HHT2-HA-TEVsite-MYC, *bar1*, *rtt109::NAT* |
| YGY675 | MATa his3Δ1 leu2Δ0 LYS2 met15Δ0 ura3Δ0 HTB2-TEV, HHT1-HA-TEVsite-MYC, HHT2-HA-TEVsite-MYC, *bar1*, *rtt109::NAT* |
| YGY721 | MATa his3Δ1 leu2Δ0 LYS2 met15Δ0 ura3Δ0 HTB2-TEV, HHT1-HA-TEVsite-MYC, HHT2-HA-TEVsite-MYC, *bar1*, *vps75::HYG* |
| YGY722 | MATa his3Δ1 leu2Δ0 LYS2 met15Δ0 ura3Δ0 HTB2-TEV, HHT1-HA-TEVsite-MYC, HHT2-HA-TEVsite-MYC, *bar1*, *vps75::HYG* |

### Growth conditions

All experiments were carried out in YPD media. G1 synchronization was achieved by incubating 700 ml of exponentially growing (OD_600_ 0.2) *bar1* cells with a final concentration of 5 ng/ml of alpha-factor (GenScript RP01002) for 3 hours. The G1-arrested cells (verified by microscopy) were centrifuged (Sorval SLA-1500, 5 minutes at 4000g, room temperature) in 3×250 ml bottles and resuspended in 100 ml prewarmed 30°C YPD supplemented with 50 ug/ml Pronase (Merck 10165921001), at which point the count-up for the time course was initiated. The 2×50 ml tubes were centrifuged (1 minute, 4000g at room temperature) and resuspended in prewarmed 30°C 50 ml YPD supplemented with 50 ug/ml Pronase and 0.2M Hydroxyurea (Bio Basic HB0528). Cells were pelleted again, resuspended in 550 ml 30°C prewarmed YPD supplemented with 50 ug/ml Pronase and 0.2M Hydroxyurea and grown shaking at 30°C for the duration of the time course. 50 ml were taken for each time point: asynchronous (taken aside before alpha-factor was added), alpha-factor arrested (before release) and 15/30/40/50/70/90/110/130/150/180 following release from alpha factor. Each 50 ml sample was crosslinked for 5 min at room temperature (RT) in 1% Formaldehyde (37% stock Baker, Cat. # 7040.1000), quenched by adding 0.125M freshly prepared Glycine (Merck G7126) at RT for 5 minutes, washed twice in 50 ml ice-cold water, pelleted and snap frozen in liquid nitrogen.

### Chromatin ImmunoPrecipitation (ChIP-seq)

ChIP was carried out as in (Yaakov et al. 2021). Cell pellets were thawed on ice and washed in 10 ml 1M sorbitol. After complete liquid removal, pellets were resuspended in 600 ul RIPA buffer (10 mM Tris pH 8.0, 140 mM NaCl, 1 mM EDTA, 0.1% SDS, 0.1% sodium deoxycholate, 1% Triton X-100, EDTA-free protease inhibitor cocktail) and transferred to chilled LoBind Eppendorf microcentrifuge tubes containing ∼500 ul of 0.5 mm zirconium oxide beads (Next Advance, ZrOB05). Cells were processed for 3 cycles in a Bullet Blender 24 (Next Advance) at level 8 for 1 min, with 1 minute on ice between cycles. Debris and lysate were transferred by piercing a hole in the bottom of the tube, placing a clean chilled tube under and centrifuging at 600xg for 5 seconds at 4C. The upper tube with the Zirconium beads was discarded and the lysate was hard spun at 17,000xg for 10 minutes at 4C. The supernatant containing cleaved solubilized 8xmyc tags was discarded, and the pellet was thoroughly resuspended in 100 ul NP buffer (10 mM Tris pH 7.4, 1 M sorbitol, 50 mM NaCl, 5 mM MgCl2, 1 mM CaCl2, and 0.075% NP-40, freshly supplemented with 1 mM β-mercaptoethanol, 500 μM spermidine, and EDTA-free protease inhibitor cocktail), and warmed to 37°C in a heatblock for 5 minutes. 100 ul NP buffer supplemented with 40 units of Micrococcal Nuclease (Worthington MNase, LS004798) were added and samples were incubated for 20 minutes at 37°C. The MNase reaction was stopped by adding 200 ul ice-cold stop buffer (220 mM NaCl, 0.2% SDS, 0.2% sodium deoxycholate, 10 mM EDTA, 2%,Triton X-100, EDTA-free protease inhibitor cocktail.), vortexing and placing on ice. Following a 30-minute ice incubation, the MNAse-treated lysates were centrifuged at 17,000xg for 10 minutes at 4C.

The 400 ul supernatant was divided into 4 separate wells of a 96-well LoBind Eppendorf plate as follows: 110 ul lysate for anti-HA IP (added 10 ul of 12CA5 hybridoma supernatant), 110 ul lysate for anti-myc (added 10 ul of 9E10 hybridoma supernatant), 70 ul lysate for anti-H3K9ac (added 5ug of ab4441) and 70 ul for anti H3K56ac (added 10 ul of Rabbit polyclonal anti-H3K56ac kindly provided by Alain Verreault (Masumoto et al. 2005)). Sequencing libraries were prepared as in (Yaakov et al. 2021). Full time courses were repeated for all strains: wildtype cells n=4 for myc & HA, n=2 for H3K9ac and H3K56ac. *rtt109* cells n=3 for myc & HA. *vps75* cells n=2 for myc, HA, H3K9ac and H3K56ac.

### ChIP-seq data processing

After demultiplexing, paired-end reads (read1 51 bp, read2 25 bp with NextSeq or 51 bp with NovaSeq) were aligned to Saccharomyces cerevisiae genome (R64-1-1) using bowtie2 (Langmead and Salzberg 2012) with the following parameters: ‘-p8 --local --very-sensitive’. Aligned reads with sizes 100–170 bases were then selected and used to calculate the genome coverage using bedtools (Quinlan and Hall 2010) with the ‘-pc’ option. Coverage files were imported into MATLAB, and normalized to a total coverage of ∼12 Mio across the entire nuclear genome without the ribosomal locus. This normalized coverage was divided into 500bp bins (or 1 kb for Figure 1E and similiar) and the mean coverage in each bin calculated. Approximately 3 million aligned genomic reads were obtained per ChIP sample (either HA, myc, K56ac or K9ac).

### ChIP-seq data analysis

#### ORI specific analysis (Figure 1C and similiar)

For each time point, the coverage of every 500bp-bin was log2-normalized to its G1 coverage using “robustfit”. Next, all 358 ORIs in the yeast genome were grouped according to their Time of Replication (ToR) into 5 ORI clusters. For each ORI cluster, the mean G1-normalized coverage across all ORIs at different distances (from 0 to 30 kb) is calculated and smoothed using robust local quadratic regression (rloess) over a 7-bin sliding window along the genome. This smoothened average is shown in Figure 1C and analogous. To calculate the correlation with DNA amount or DNA replication speed (Figure 1D and analogous) at the ORI cluster with the earliest ToR for each time point, we first calculated the DNA profile by fitting a 5-parameters logistic curve to the mean G1-normalized HA profile. As DNA replication proceeds away from ORIs, the DNA replication rate is proportional to the first derivative of this fit along the distance. For each time point, Pearson’s correlation between the mean G1-normalized profiles (HA, myc, K56ac and K9ac) around the earliest replication ORI cluster and the fitted DNA profile and replication rate was calculated and shown in Figure 1D.

#### Genome-wide analyses (Figures 2C, 2D, S2B and analogus)

For genome-wide analysis, the 500-bp bins were clustered into 96 replication clusters first by their ToR (48 groups), and then by their replication dynamics in HU (2 clusters per group = 96 clusters). For each sample, we determined the median coverage (not normalized by G1) and ToR in each cluster. As the coverage is relative and does not account for the increase in total DNA/HA (nucleosome) during S phase, we next calculated the increase in total HA, based on the observation that DNA /HA in each cluster increase concomitantly during S-phase (Bar-Ziv et al. 2016; Frenkel et al. 2021). For two consecutive time points, the increase in total HA was set so that the absolute HA level of slowest replicating cluster (either the one with the highest ToR or the lowest ToR, for early and late time point respectively), with the strongest decrease in relative coverage, stays constant once the change in total HA is accounted for. Next, the cumulated HA increase from time 0 up to a given time point was taken as the total HA level for this time point (raising from 1 to ∼1.4 during the time courses). Finally, for each time point, the absolute HA level in each cluster results from multiplying a cluster’s relative HA level with its total HA, and the replicated fraction (Figure 2C) by subtracting 1. The replication rate in cluster i at time point t, r(i,t), is calculated as:

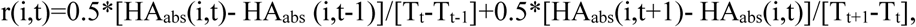

where HA_abs_ is the absolute HA level in cluster i at time point t and T_t_ is the time since HU release at time point t.

For Figure 2C, the ratio median myc over median HA level (not absolute) of each cluster at every time point is compared against its current replication rate or replicated fraction (Pearson’s correlation of all time courses is summarized in Figure S2B). The fork-dependent turnover (Figure 2C, D) is calculated as the slope of the robust linear fit (robustfit) between the myc-ha-ratio and the replication rate. The fork-dependent turnover across all time points is summarized in Figure 2D and similar.

#### Time delay analysis (Figure 1E)

To calculate the time delay between the dynamics of different epitopes (HA vs. myc/K9ac/K56ac), the genome was divided into 1kb-bins and the mean coverage in each bin was calculated. For every sample in a time course, the epitope level of every bin was log2-normalized against the corresponding level in alpha factor arrest (G1) using robust fit, and the resulting level smoothened along the genome using local robust linear regression with an 11-bin window. Afterwards, the G1-normalized dynamics across the time course were fitted with a smoothing spline and the fit used to interpolate the change in 5 minute intervals. Next, we correlated the change across all bins in HA level at each interpolated time point with the change in the other epitopes at every other time point (n^^2^ correlations for n interpolated time points). The correlation for each delay and epitope, tau, is calculated as the mean correlation, Cr, between all time point pairs separated by a certain delay:

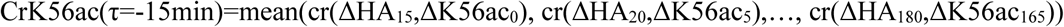

Where cr(x,x) is Pearson’s correlation between two samples and ΔXX_t_ is the interpolated change at of epitope XX at time point t.

To account for inherently different noise levels between the epitopes, all mean correlations of one epitope were normalized to its maximal correlation, i.e. correlation score.

#### Nucleosome-specific processing (Figures 3D, S2C, S5A, S5B)

Nucleosome were analyzed as described in (Yaakov et al. 2021). In brief, genome-wide coverage was normalized to a total coverage of 10^^8^, and the mean normalized coverage 100-bp around each nucleosome position (+/- 50bp) was calculated.

Log_2_ turnover (myc/HA) and epitope enrichment (K56/HA), EnrX, for each nucleosome (Figure S2C) was calculated as: EnrX = log2(X(i,t)+1) – log2(HA(i,t)+1), where X(i,t) is the level of epitope X on nucleosome i at time point t.

Late replicating and late replicating and highly turning over nucleosomes (Figure 3D) were selected based on the ToR cluster with less than 10% replication at the last time point across all time course (mean) and a mean turnover (log2(myc/ha)) greater than 1.5 in G1 arrested cells, respectively.

For HA/H3 level by ToR analysis (Figure S5), each nucleosome was assigned a ToR based on its position and the mean epitope level across all nucleosomes with a particular ToR (+/- 1 min) calculated and shown.

## Data and Code availability

All Next Generation Sequencing (NGS) data generated in this project is available on GEO (GSE: 193044). Additional data used to compare double- and single-tagged exchange sensor (Fig S5) is also available on GEO (GSE: GSE157402). All MATLAB scripts to analyze the data and generate the figures can be found on GitHub (https://github.com/barkailab/Yaakov2022).

## Acknowledgements

We thank Y. Voichek and N. Frenkel for comments on the manuscript. We thank our lab members for discussions throughout. We are very grateful to Alain Verreault for the H3K56ac antibody. This work was funded by the Israel Science Foundation FIRST Program (grant No. 966/19), ERC and the Minerva Foundation.

## Author contributions

F.J. performed all analyses. G.Y performed all experiments. All authors designed the experiments, discussed the results and wrote the paper.

